# Characterization of the Lipidome of Neurons in Mouse Brain Nuclei using Imaging Mass Spectrometry

**DOI:** 10.1101/2025.09.04.674184

**Authors:** Cristina Huergo, Laura De las Heras-García, Jone Razquin, Yuri Rueda, Cristina Miguélez, José A Fernández

## Abstract

Understanding the molecular composition of the brain at cellular level is essential for deciphering the metabolic alterations associated with brain diseases. Furthermore, the different prevalence of some neurological diseases between males and females highlight the importance of incorporating gender factor in such studies. Here, we demonstrate that using imaging mass spectrometry in negative polarity it is possible to isolate and characterize the lipidome of specific neuronal populations in the mouse brain, including the locus coeruleus (LC), mesencephalic neurons and the substantia nigra pars compacta (SNc). Neuronal identity was validated through immunofluorescence on adjacent serial sections. Comparative analysis revealed that each neuronal population presents a distinct and well-defined lipidic profile, with differences extending across all lipid classes analyzed. Regarding sex-based differences, we found discrete differences in phosphatidylcholine/phosphatidylethanolamine-ether, phosphatidylinositol and sphingomyelin LC neurons. Lipidomic differences were more pronounced in mesencephalic neurons, whereas no significant sex-defendant differences were observed in SNc lipid composition. These findings lay the groundwork for future studies aimed at identifying lipid metabolic dysregulations in the context of neurodegenerative diseases.

## INTRODUCTION

The introduction of imaging mass spectrometry (IMS) for the study of lipids has revealed a complex landscape, in which each cell type displays a well-defined lipidome. Histological structures previously considered homogeneous now show a rich diversity of lipid fingerprints. For example, recent studies have uncovered variations in lipid expression across different areas of the nephron, potentially correlating with the relative abundance of specific cell populations.^[1-3]^ The retina is another beautiful example of complexity from a lipidomic perspective.^[4]^ Furthermore, investigations into the human colon lipidome have shown that not only distinct cell populations but also colonocytes at varying stages of maturation exhibit a gradient of lipid profiles.^[5-7]^

Among all tissues, the brain presents the highest level of complexity. Each brain nucleus performs highly specialized functions and, therefore, displays a distinct cell composition.^[8-14]^ Not only neurons but also glial cells in the central nervous system (CNS) are highly specialized, with morphology and function varying between brain areas.^[9, 15]^ Consequently, considerable efforts are underway to map brain cell populations as a first step toward understanding brain function. Such efforts include several techniques to obtain a multimodal view: optical microscopy, MRI, cell-resolution transcriptomics etc. Within this context, IMS has gradually emerged as a valuable component of this multi-modal approach, owing to advances in spatial resolution and sensitivity. From the first images recorded at a very modest spatial resolution,^[16-18]^ substantial progress has been made toward capturing increasingly detailed views of brain regions. ^[19]^

To fully understand brain pathologies, however, achieving cellular resolution is essential. Different cell types exhibit distinct lipid profiles; for example, astrocytes are enriched in phosphatidylinositol (PI), microglia show higher concentrations of sphingomyelin (SM) and phosphatidylglycerol (PG), whereas neurons are particularly enriched in phosphatidylethanolamine (PE) and phosphatidylcholine (PC).^[20]^ Therefore, developing methodologies with sufficient spatial resolution to isolate cell-specific lipidomes or well-defined cell populations is a critical goal.

Here, we focus on the neuronal lipidomic profile. We present a mass spectrometry imaging study of the lipidome of three types of neurons in mouse brain: noradrenergic neurons in the locus coeruleus (LC), the sensory neurons in the mesencephalic trigeminal nucleus (Me5) and the dopaminergic neurons of the substantia nigra pars compacta (SNc), carried out at 10 μm/pixel of spatial resolution. This resolution permits not only the identification of individual neurons, but also the spatial separation of their soma from projecting processes.

The LC, although small and densely packed, is the principal source of noradrenaline in the CNS, and plays a key role in functions such as arousal, sleep and stress responses.^[21, 22]^ Me5 neurons are primary sensory neurons responsible for proprioception of the masticatory muscles, projecting to central branches into the trigeminal motor nucleus.^[23]^ Compared to LC neurons, these are substantially bigger and are found isolated from one another, which largely facilitates segregation of their lipid profile from the surroundings. Dopaminergic neurons of the SNc form a compact band of small cells that can be readily distinguished from neighboring structures. This nucleus plays a crucial role in motivation, memory, and motor control. ^[24, 25]^

Isolating the lipid signature of the three types of neurons allowed us to identify differences in composition, providing a first step toward characterizing changes in their lipid profile in the context of several pathologies. This approach is particularly relevant for neurodegenerative diseases such as Parkinson’s disease (PD), in which monoaminergic neurons, both noradrenergic in the LC and dopaminergic in SNc, are especially vulnerable and undergo early degeneration.^[26]^ Some of the characteristics that make these two neuromelanin-containing nuclei more susceptible to degenerations may include their highly branded unmyelinated axons, continuous pacemaking activity, elevated cytosolic Ca^2+^ concentrations and high levels of mitochondrial oxidative stress. ^[27-29]^ In addition, an altered lipid profile could serve as an indicator of increased risk for PD. ^[30]^ A comprehensive understanding of the neuronal lipidome and the potential metabolic alterations that occur during the disease is essential for elucidating its underlying mechanisms and identifying new therapeutic targets. Furthermore, the inclusion of both male and female animals in the study permits the exploration of potential sex-related differences in lipid composition, which may contribute to understanding sex-specific vulnerabilities to neurological disorders.

## METHODS

### Chemicals

To perform the experiments, the following chemicals were used: Acetone (Panreac, Spain, 99.5%), Isofluorane (Nuzoa, Spain), Isopentane (Sigma-Aldrich, Germany, ≥99% purity), Methanol (Sigma-Aldrich, Germany, 99.8%), Mowiol 4-88 (Sigma-Aldrich, Germany, >99.5% purity), NaCl (Sigma-Aldrich, Germany, ≥99.5%), NaHPO_4_ (PanReac, Spain, ≥99% purity), NaH_2_PO_4_·H_2_O (PanReac, Spain, ≥99% purity), Paraformaldehyde (Sigma-Aldrich Germany, 95.0-100.5%), Tissue-Tek® O.C.T. Compound (Sakura Finetek, USA), Triton™ X-100 (Sigma-Aldrich, Germany, <1.00% water (Karl Fischer)).

### Animals

Experiments were performed on 18 adult (12 weeks old, weight 20-30g) male and female C57BL/6J mice (Envigo, Spain), group housed with no more than 5 animals per cage, given access to food and water ad libitum, and maintained on a 12:12-hour light/dark cycle. All procedures were performed in compliance with the European Communities Council Directive on “The Protection of Animal Uses for Scientific Purposes” (210/63/EU) and Spanish Law (RD 53/2013) for the care and use of laboratory animals. Protocols were approved by the Bioethical Committee for Animal Research of the University of the Basque Country (Spain) (UPV/EHU, CEEA, M20/2021/215 and M30/2021/220).

For brain extraction, anesthesia was induced with 4% isoflurane delivered in oxygen at a flow rate of 1□L/min. Once the absence of reflexes was confirmed, animals were immediately decapitated to obtain fresh brain tissue. Brains were rapidly extracted and snap-frozen in pre-cooled isopentane (Sigma-Aldrich, Germany), cooled on dry ice to approximately −40□°C, for 30–60 seconds. Tissues were then transferred to dry ice and subsequently stored at −80□°C until further processing.

### MALDI-IMS experiments

Using a cryostat (Leica CM3050 S, Leica Biosystems), between 12 and 16 coronal sections, each 25 µm-thick, were obtained per mouse brain. The sections were mounted onto ITO-covered slides (Intellislides, Bruker Daltonics, Germany) for IMS and superfrost slides (Epredia, USA) for immunohistochemistry (IHC) and stored at -80 ºC. Those for IMS were covered with 1,5-diaminonaphthalene, using an in-house designed sublimator ^[31]^ and introduced into the MALDI (matrix-assisted laser desorption/ionization) source of the mass spectrometer (TIMS ToF Flex, Bruker Daltonics, Germany) available in the analytical services of the University of the Basque Country (UPV/EHU). Data acquisition was performed at 10 µm/pixel, using 100 laser shots per pixel and a laser energy of ∼40 μJ/pulse. Mass resolution was set at ∼60,000 at m/z = 1000. All experiments were recorded in negative-ion mode with an observation window of 300-1300 Da, where most glycerophospholipid and sphingolipid classes appear. Altogether, 62 sections from 10 animals containing Me5, of which 51 also contained LC were measured and the two sides averaged. Regarding SN, 21 sections of 12 animals containing SNc were measured. No statistically-significant differences between the lipid signature of the neurons of each nucleus between sections of the same animal were found.

The data obtained in this way were processed using SCILS (Bruker Daltonics, Germany). To extract the peaks, the threshold intensity was set such that the number of peaks (m/z channels) was ∼400. Then, supervised k-means segmentation was used to extract the segment containing the neuron population of interest. The mass spectrum of the selected segment was exported as an excel file and analyzed using in-house developed software.

### UHPLC-MS experiments

To help with the identification of the lipid species detected in the IMS experiments, lipid extracts of the LC, Me5 and SNc of sixanimals were obtained following an isopropanol (IPA) method ^[32]^ and HPLC-MS lipidomic analysis was done as described somewhere. ^[3]^ Briefly, the lipid extracts were injected into an HPLC column coupled to a QExactive HF-X (ThermoFisher) mass spectrometer. The analysis was performed in positive- and negative-ion modes, after optimizing the parameters using the Splash LipidoMix (Avanti Polar Lipids, Alabaster, AL) standard. MS data were acquired and processed using the Xcalibur 4.1 package, with 5 ppm tolerance for lipid precursor and fragment ions. Assignment of the lipid species was carried out using LipidSearch software version 5.1.

### Immunofluorescence

Attempts to record IMS and immunochemistry (IHC) experiments in the same section resulted in sub-optimal fluorescence images. Therefore, serial sections were used for IHC so the true nature of the neurons could be certified. The IHC experiments were carried out following the protocol in ref [21]. First, sections stored at −80□°C on microscope slides were thawed and dried at room temperature in desiccation boxes with silica gel beads. Then, they were fixed with 4 % paraformaldehyde in phosphate buffered saline (PBS 0.1 M, pH□=□7.4) for 15 min at room temperature and subsequently washed with PBS (3 times, 5 min). The first permeabilization was done with cold methanol:acetone (1:1) at −20□°C for 10 minutes. After washing with PBS, tissue sections were permeabilized again and non-specific labelling was blocked using 5% of normal goat serum (NGS, Sigma-Aldrich, Germany) in 0.1 M PBS and 0.5% Triton X-100 (Sigma-Aldrich, Germany) for 2 hours at room temperature. Next, the samples were incubated with primary antibodies overnight at 4°C in 5% NGS and 0.5% Triton X-100. After washing with PBS, the corresponding fluorochrome-conjugated antibodies Alexa Fluor 488 or 555 conjugated goat secondary antibodies (1:400; Invitrogen) were applied and the samples were incubated during 2 hours at room temperature. After washing with PBS, they were incubated with Hoechst 33342 (1:10000; Invitrogen) for 10 min at room temperature for nuclei labelling and washed again. They were mounted using Mowiol 4-88 mounting medium (Sigma-Aldrich, Germany). Primary antibodies used for immunofluorescence (IF) on these tissues included: chicken anti−tyrosine hydroxylase (TH, 1:2000; #ab76442 Abcam) for labeling noradrenergic and dopaminergic neurons in the Lc and SNc, respectively, and rabbit anti-p92 (1:500; #ab72210 Abcam) for Me5 neurons.

All IF images were acquired as z-tack images using the Zeiss LSM800 confocal microscope (Plan Apochromat 10x NA: 0,45 d:2.1mm; Plan Apochromat 20xAir NA: 0,8 d:0,55mm DIC) with the same settings for all samples. Fluorescence image processing was performed with the ImageJ software (National Institutes of Health; NIH).

### Statistical analysis

Statistical analysis was carried out using SPSS Statistics 18.3 (IBM, Armonk, NY, USA). The statistical tests used were Levene test, ANOVA univariate statistical analysis and Tukey/Games Howell post hoc analysis. The Levene test determines the homogeneity of variance (H_0_ = groups have equivalent variance), to choose the post hoc method: we used Tukey if Levene test p ≥ 0.05 and Games Howell if Levene test p ≤ 0.05. PCA analyses were carried out using Orange Biolab V.2.7.8 (Ljubljana, Slovenia).

## RESULTS

As illustrated in **Figure 1**, the LC is a bilateral nucleus located in the hindbrain, beneath the cerebellum and adjacent to the fourth ventricle. It is densely populated by TH-positive neurons responsible for noradrenaline synthesis. Lateral to the LC are the Me5 neurons, which are large, isolated cells with round to oval morphology that stain for the p92 protein. The SN is anatomically divided into the dorsally located pars compacta (SNc), rich in dopaminergic TH-positive neurons, and the ventral pars reticulata (SNr), which mainly contains GABAergic neurons.

**Figure 1.**
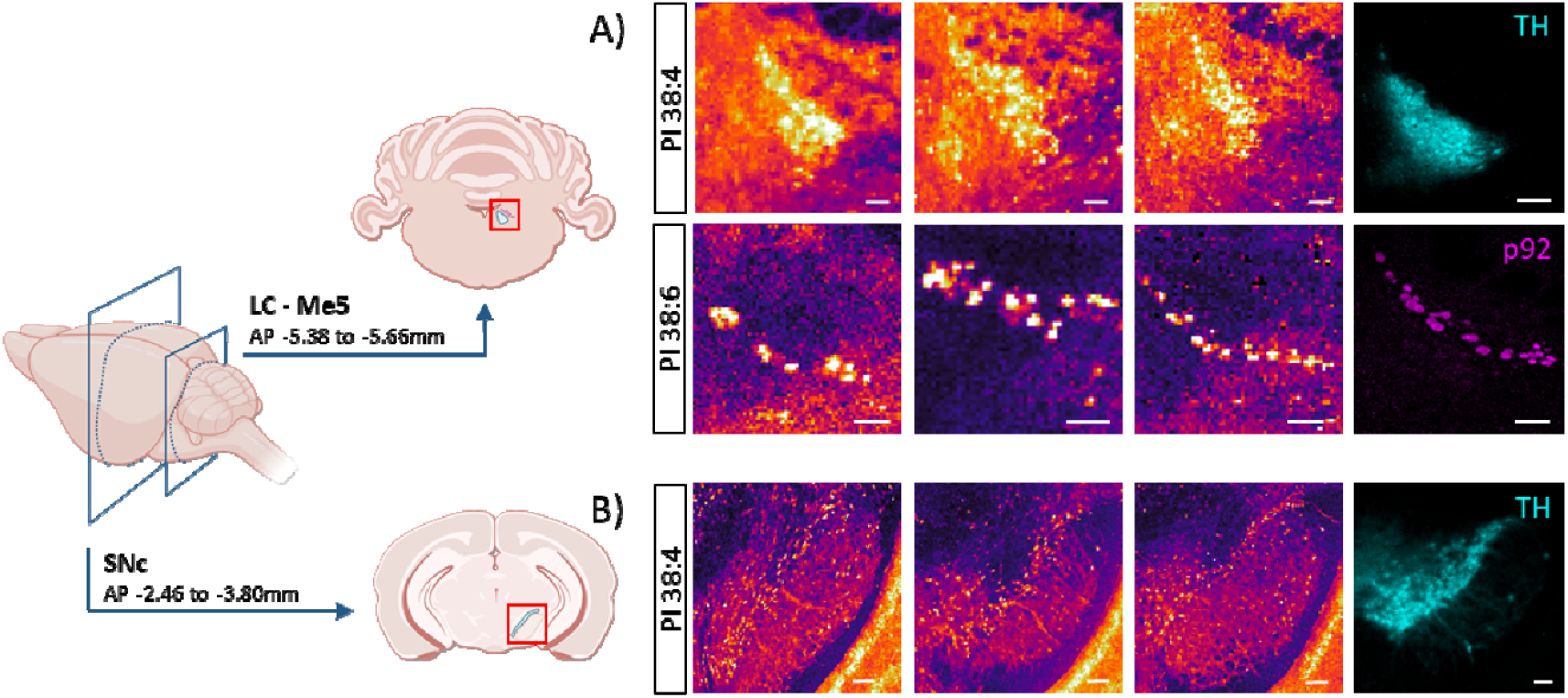
Coronal mouse brain sections measured using MALDI-IMS. Neurons of LC (upper panels) present a larger abundance of phosphatidylinositol (PI) 38:4. The distribution of that lipid species matches very well the IF images of TH (tyrosine hydroxylase). Conversely, Me5 neurons are enriched in PI 38:6 (middle panels). The distribution of this lipid reproduces faithfully the p92 IF images. The neur ns in SNc also present a large abundance of PI 38:4. Comparison with the TH image helps identifying the distribution of the neurons. Images of lipid distribution recorded at 10 μm/pixel in negative-ion mode. Scale bar 100 μm. Icons created with BioRender.com.

Representing the distribution of PI 38:4 (**Figure 1**) it is possible to visualize the LC and the SNc. The images obtained at 10 μm/pixel show that neurons are densely packed, following also the distribution observed in the TH IF image. Interestingly, the Me5 neurons, which are very close to the LC, are deficient in PI 38:4 and appear as dark spots. The opposite happens with the distribution of PI 38:6, which is more abundant in Me5 neurons.

Segmentation of the images is not an easy task. The segmentation algorithm can segregate the neuron bodies of the LC from their processes (**Figure 2**). However, it is not able to group all the pixels of the Me5 into a single segment. Conversely, it creates several groups and segregates the nuclei of the neurons from the rest of the neuronal body. Therefore, manual selection of the neurons was done, using the p92 IF image and the PI 38:6 distribution as a guide. Neurons in the SNc are relatively small, their bodies occupy 1-4 pixels and, therefore, they can only be isolated at high spatial resolution. Close to the SNc lies the SNr, which contains two distinct neuronal subpopulations: parvalbumin-positive and parvalbumin-negative neurons. ^[33]^ Attempts to isolate those neuronal subpopulations were unsuccessful. While the segmentation algorithm was able to distinguish the neuron’s surrounding environment, it failed to accurately separate the neuronal bodies themselves. Therefore, we limited the study to the lipidomes of LC, Me5 and SNc neurons.

**Figure 2.**
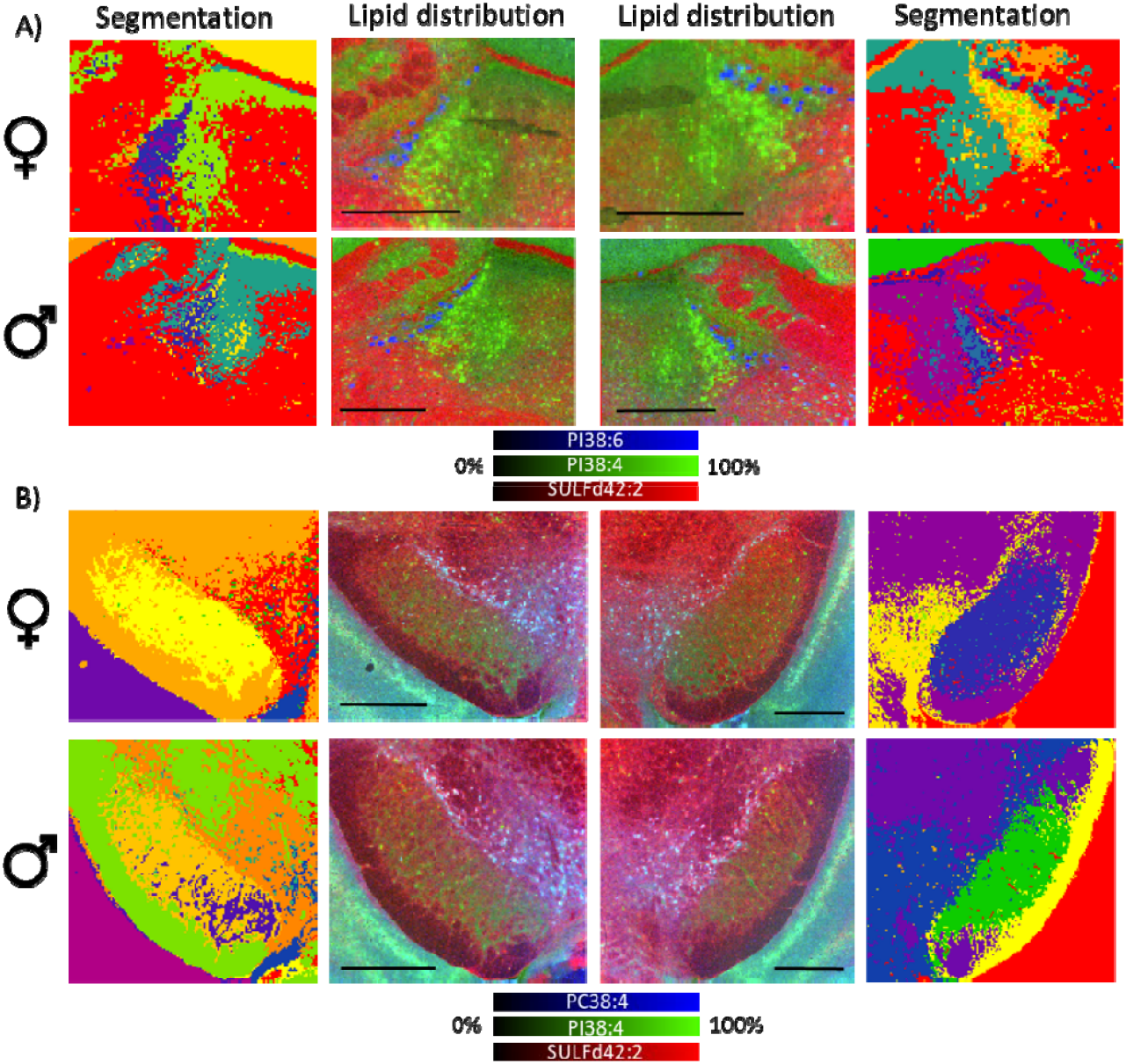
Comparison between the segmentation (first and last columns) and the distribution of three lipids (central columns) in se tions containing A) LC and Me5 (upper two rows) and B) SNc (lower two rows). LC neurons appear as bright green in the central images, while Me5 neurons appear as blue pixels. In the segmentation images, LC is clearly visible, while the Me5 neurons are difficult to locate, making it necessary to manually choose the pixels containing them. In the images of SN, the SNc neurons appear as blue spots and are also present as individual pixels in the segmentation images. Experiments recorded in negative-ion mode at 10 μm/pixel. Scale bar 500 μm.

Analysis of the lipid fingerprint of the three brain nuclei demonstrates that in all of them PC/PE are the most abundant lipid families, followed by PI and PC-ether/PE-ether (PEe/PCe). However, there are important differences in the relative abundance of those families (**Figure 3A** and **Figures S2-S4** of the ESI). Both the statistical analysis and the PCA indicate that the neurons in each nucleus exhibit a distinct and well-defined lipidome, clearly differentiable from the others. LC neurons appear to be more enriched in SM, PI, and PEe/PCe. In contrast, Me5 neurons display higher levels of PE and PC, whereas SNc neurons are enriched in phosphatidylserine (PS).

**Figure 3.**
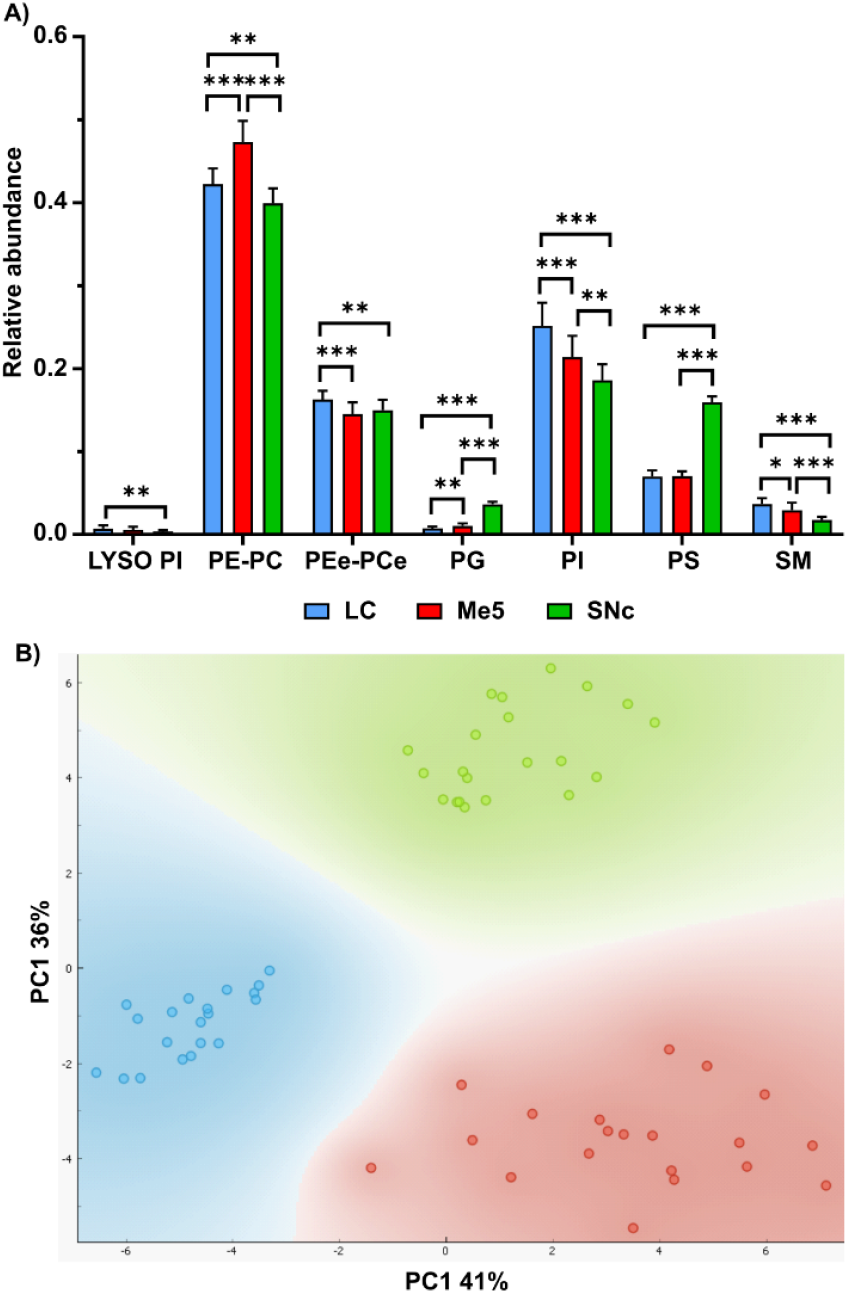
(A) Comparison of the relative abundance of the main lipid families studied in this work between LC, Me5 and SNc neurons; (B) principal components analysis (PCA) of the lipid fingerprints of the three types of neurons. * = p<0.05, ** = p<0.01, *** = p<0.001.

Comparison between the neuron profiles in males and females (**Figure 4** and **Figures S5-S10** of the ESI) shows consistent differences in lipid composition between sexes in LC and Me5. However, the PCA analysis does not show a clear separation in SNc samples. Statistical analysis of the relative abundances of the lipid classes studied here also demonstrate the existence of statistically significant differences in LC and Me5 but not in SNc neuron lipid composition between male and female.

**Figure 4.**
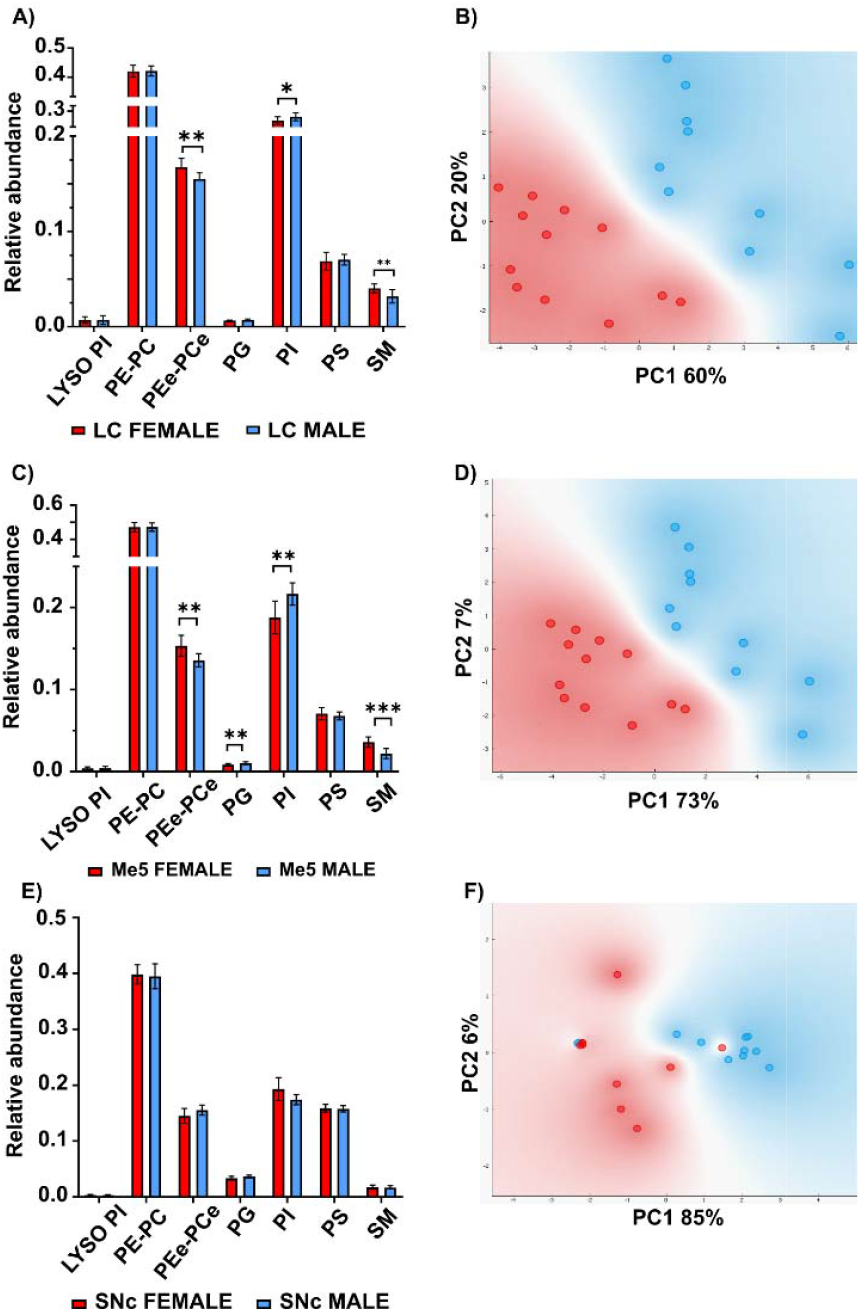
Comparison between the lipid composition of LC (A), Me5 (C) and SNc (E) neurons of male and female mice. The corresponding PCA analysis is shown in panels (B), (D) and (F). * = p<0.05, ** = p<0.01, *** = p<0.001.

## DISCUSSION

### Lipidomic Signatures of Distinct Neuronal Populations

Cell diversity reflects the richness and complexity of the brain, with different neuronal nuclei exhibiting distinct morphologies, functions, and metabolic profiles, which in turn influence their vulnerability to neurodegeneration. In this context, MALDI-IMS experiments performed at 10 μm/pixel allowed us to isolate the lipid composition of different neuron populations, such as those in the LC and SNc, regions that possess unique metabolic characteristics and degenerate in PD, in contrast to other neurons like those in the Me5.

To the best of our knowledge, this is the first study in which lipid profiles of specific neuronal populations have been directly extracted from brain sections. Previous works, such as that performed on Purkinje cells, identified only a limited number of lipid species, as the primary aim was to demonstrate the technical feasibility of MALDI-IMS combined with histological analysis ^[34]^. Other studies have reported neuronal lipid profiles using either cell cultures ^[35-37]^ or complex procedures for isolating and sorting individual neurons.^[13, 20, 38-42]^ For example, Merrill et al ^[43]^ identified more than 40 lipids from dentate gyrus granulate cells and CA1 pyramidal neurons of the hippocampus using patch clamp to extract single neurons from the tissue, followed by nanoflow liquid chromatography/high-resolution time-of-flight mass spectrometry (nLCMS) analysis. They did not report significant differences in the lipid composition of these neurons. However, they observed substantial differences in lipid composition between neurons and their surrounding tissue.

Our results generally align with those of Merrill et al ^[43]^ in terms of the lipid species detected, but several key differences were noted. For example, the most abundant PC species in their study was PC 32:0, whereas in our work PC 34:1 predominates (**Figure S2**). Our observation is consistent with reports from studies on single HT22 neurons, ^[44]^ SNCA-A53T neurons ^[45]^ or neurons sorted by flow cytometry. ^[20]^ Neumann et al also reported a PC 32:0/34:1 ratio >1 in neurofilament light chain (NfL)-positive cells from cultured rat cerebellum.^[35]^ Methodological differences may account for these discrepancies: Merrill et al. used positive polarity while our experiments were conducted using negative polarity; our m/z signal includes overlapping contributions from PE 36:1 and PC 34:1; and our measurements focused on neuronal somas, while their analysis encompassed the entire neuron, including dendrites and axon. However, the most likely explanation is that each neuron population possesses a unique lipidomic fingerprint reflecting its specific function and metabolic profile.

With respect to PI, the authors of ref [43] reported only two species: PI 38:4 and 38:5, with the latter being more abundant. While these were also the most abundant PI species in our study (Figure S3), their relative intensities were reversed. Fitzner et al^[20]^ also identified PI 38:4 as the most abundant species, followed by PI 36:4, and noted substantial regional differences in their relative abundance. Notably, LC neurons in our data show substantially higher PI 38:5 levels compared to Me5 and SNc neurons. Therefore, comparison with the results in refs [20, 35, 43-45] reinforces the hypothesis of a high specificity of lipid composition in each neuron type. PI is a very specific lipid of neurons and astrocytes, although certain species are more abundant in neurons, as can be seen in **Figure 1**. The presence of PI in neurons is restricted to the membranes where it is produced or localized to the inner leaflet of the lipid membrane. The myo-inositol head group can be selectively phosphorylated to produce up to seven different species that play very specialized roles, ^[46]^ such as regulation of calcium concentration or modulating the activity of specific proteins via direct binding. It is therefore no surprise that such specialized lipids exhibit different molecular compositions in distinct neuron populations.

Differences in PS were more modest across neuronal types. Consistent with prior studies,^[20]^ PS 18:0/22:6 was the most abundant species. PS is also localized in the inner leaflet of the plasma membrane together with docosahexaenoic acid (DHA; FA 22:6), regulating Akt activity, which is crucial for neuronal survival. ^[47-51]^ DHA accumulates mostly in PS and PE, also in good agreement with the data reported here. PS is involved in various cerebral functions, including membrane signaling, neuroinflammation, neurotransmission, and synaptic remodeling.^[50]^

PG are phospholipids very specific to mitochondria. They are usually located in the inner mitochondria membrane, where they are used to produce cardiolipins. ^[52]^ PG are usually present in very low abundance, but their alteration may be associated with certain neurological diseases. ^[53]^ We observed increased PG levels in SNc neurons, particularly in species enriched in polyunsaturated fatty acids (PUFA) such as arachidonic acid (AA) and DHA. Examination of the AA and DHA content in all three neuronal types (Figure 5) shows large variations between the nuclei, with a larger abundance of 22:6 in Me5 and an increased abundance of 20:4 in the neuros of the two nuclei vulnerable to PD.

**Figure 5.**
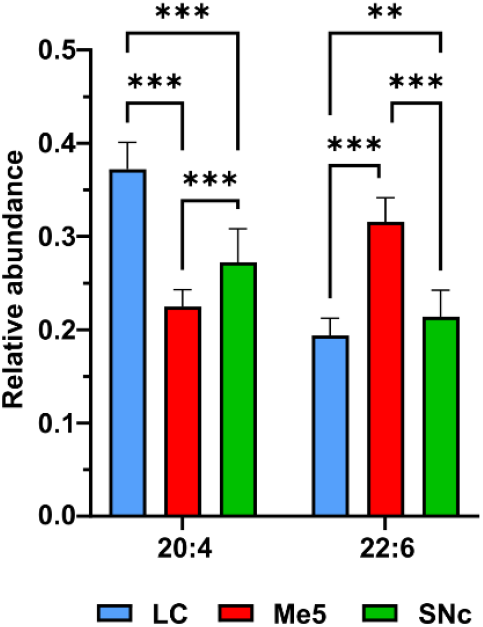
Relative abundance of lipid species containing AA and DHA in neurons of LC, Me5 and SNc. Calculation was carried out by adding the relative intensities of all the species containing the mentioned fatty acids.

Regarding SM, the molecular species strongly contrast with those from PG, as they do not contain PUFA. The most abundant species are SM 36:1, 36:2, 38:1 and 42:2, in good agreement with previous publications.^[20, 43, 44]^ The data presented here point to strong variations in the relative abundance of most of the species detected. For example, there is an almost complete absence of SM 34:1 and 38:1 in SNc neurons. Sphingolipids are known for their role as modulators of membrane fluidity and the function of some receptors and proteins. ^[54-57]^ They are enriched in saturated fatty acids, which pack more tightly in the presence of cholesterol. Therefore, they are able to form ordered domains in the lipid membrane, facilitating proper protein localization. ^[56, 58]^ The marked differences in SM levels between the three types of neurons may be related to the differences in receptor expression. Actually, the list of ion channels and signaling receptors in the CNS that need to be anchored to a membrane microdomain and therefore, are regulated by SM membrane composition is continuously expanding. ^[56]^

Although LC neurons share morphological similarities with those in the SNc, and both degenerate in PD, this degeneration seems to not occur simultaneously. Evidence suggests that the LC is affected earlier than the SNc, and that the loss of LC neurons may influence or predispose the subsequent SNc degeneration.^[59]^ Their distinct lipidomic fingerprints, indicative of significant metabolic differences, may be linked to this temporal sequence and could help explain their differential vulnerability to neurodegenerative processes such as PD. ^[60]^

### Sex Differences in Neuronal Lipid Composition

The results presented in **Figures 4** and **S5-S10** demonstrate the existence of differences in lipid composition between male and female neurons. However, these differences are neuron-type and lipid-class dependent. In LC neurons, differences between males and females were observed only in PCe/PEe, PI and SM. Across individual species, a general trend toward lower abundance in males was observed. These findings are in line with existing literature reporting sex differences in LC neurons at anatomical, electrophysiological, and molecular levels. Females have been shown to exhibit a greater number of neurons, more elaborate dendritic arborization, and distinct electrophysiological and molecular characteristics. ^[61-64]^

Interestingly, no differences were observed in PC/PE species, confirming the decoupling of both lipid classes. PC/PE are the main components of the lipid membranes and play mostly a structural role, ^[65]^ although they may also be involved in more specialized tasks. ^[66]^ In contrast, the differences in PCe/PEe are not restricted to a group of species with a certain fatty acid composition. Conversely, approximately half of the detected species present lower abundance in males. Plasmalogens are known to have multiple functions, from modulating membrane properties to acting as antioxidants or metal ion chelators. ^[67]^

Male Me5 neurons also show a reduction in PCe/PEe, but in this case, the reduction is mostly limited to PUFA-containing species. Conversely, male Me5 neurons have a higher abundance of DHA-containing species and a decrease of AA-containing species. DHA and AA are signaling molecules with opposite effects: while DHA is involved in the production of resolvins, anti-inflammatory lipids, AA is used to produce prostaglandins and to trigger the inflammatory response.

Sex differences were also found in PI and SM content. In LC and SNc neurons, males had modest increases in AA-containing PI species. Conversely, SM species were generally lower in male neurons. The changes observed in the LC contrast with those in the SNc. Notably, in SNc neurons, only one PI species (PI 34:0) showed a statistically-significant sex-related difference.

It is difficult to explain all the observed differences in lipid expression in neurons between male and female mice, as many factors may be considered. Previous studies using HPLC-MS also reported sex-dependent differences in brain lipid composition and even distinct alterations in response to diet. ^[68, 69]^ Moreover, estrogens are known to regulate lipid homeostasis in mice. ^[70]^ Another intriguing finding is that sex-related differences in lipid composition were robust in the LC and Me5, but absent or very mild in the SNc. This observation is consistent with previous studies showing that the LC is a clearly sexually dimorphic nucleus, ^[61]^ whereas the SNc, less thoroughly investigated in this regard, does not exhibit such pronounced dimorphism.

The sex-related differences reported here may contribute to understand the known sex disparities in the incidence and progression of several neurological diseases. Altered lipid profiles have been implicated in various CNS disorders, including neurodegenerative diseases. ^[71-73]^ Although direct evidence of sex differences in human neuronal lipid composition is currently lacking, some early studies suggest a role for lipid signaling molecules, such as lysophosphatidic acid, in neuropsychiatric and neurodegenerative disease pathogenesis.

^[74]^ Moreover, the Alzheimer’s disease mouse model *Abca7* knockout displayed sex-specific lipid dysregulation in the brain. ^[75]^ These findings underscore the need to investigate whether lipidomic sex differences play a role in human disease susceptibility and progression.

## CONCLUSIONS

Although the general composition of brain lipids has been previously described in both mice and humans, ^[20, 41, 76-79]^ to our knowledge, this is the first study that isolates and analyzes lipid profiles of neurons in specific nuclei like LC, Me5 and SNc, directly from tissue sections. The results demonstrate that each neuronal population exhibits a distinct lipidomic signature, highlighting region-specific roles of lipids in neuronal function and vulnerability. In addition, sex-related differences were identified in the LC and Me5, supporting the hypothesis that neuronal lipid composition may contribute to sex-specific susceptibility to neurological disorders. These findings establish a foundation for future research aimed at elucidating how lipid diversity influences neuronal physiology and disease pathogenesis.

## ASSOCIATED CONTENT

### Supporting Information

The Supporting Information is available free of charge on the ACS Publications website.

Supporting Information-1: additional figures with the comparison of lipid distribution between neuron types and male/female animals in PDF format

Supporting Information-2: table with the lipids detected and acyl composition for those also identified by UHPLC-MS in XLSX format

## AUTHOR INFORMATION

### Present Addresses

†If an author’s address is different than the one given in the affiliation line, this information may be included here.

### Author Contributions

The manuscript was written through contributions of all authors. / All authors have given approval to the final version of the manuscript. / ‡These authors contributed equally.

## ACKNOWLEDGMENT

The authors thank for technical and human support provided by SGIker (UPV/EHU/ERDF, EU) Core Facility for Analytical and High-Resolution Microscopy in Biomedicine, and Bizkaia General Animal Facility.

This work was supported by grants PID2021-126434OB-I00 and PID2021-127918NB-I00 funded by MCIN/AEI/10.13039/501100011033 and ERDF A way of making Europe. It has also been funded by the Basque Government (IT1706-22 and IT1491-22). This research was conducted in the scope of the Transborder Joint Laboratory (LTC) “non-motor Comorbidities in Parkinson’s Disease (CoMorPD)”. LDH and JR hold PhD grants from the University of the Basque Country.

## Insert Table of Contents artwork here

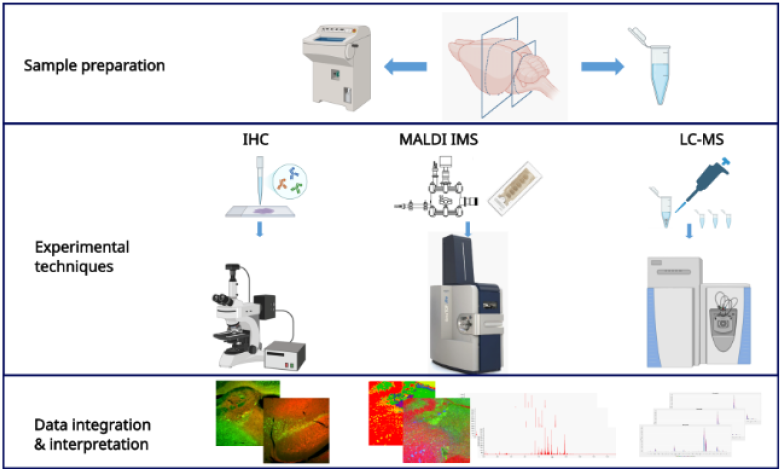

## Notes

### Competing Interest Statement

The authors have declared no competing interest.

