## Supplemental information for "Characterization of the Lipidome of Neurons in Mouse Brain Nuclei using Imaging Mass Spectrometry"

[1]Dep. of Physical Chemistry, Fac. of Science and Technology, University of the Basque Country (UPV/EHU), Barrio Sarriena S/N, 48940 Leioa, Spain; [2]Dep of Pharmacology, Faculty of Medicine, University of the Basque Country (UPV/EHU), Barrio Sarriena S/N, 48940 Leioa, Spain; [3] Neurodegenerative Diseases, Biobizkaia Health Research Institute, Barakaldo, 48903, Spain; [4]Univ. Bordeaux, CNRS, IMN, UMR 5293, F-33000, Bordeaux, France.; [5]Dep. of Neuroscience, Universidad del Pais Vasco (UPV/EHU), Barrio Sarriena S/N, 48940, Leioa, Spain;[6] Department of Physiology, Faculty of Medicine and Nursing, University of the Basque Country (UPV/EHU), B. Sarriena, s/n, Leioa 48940, Spain

Index

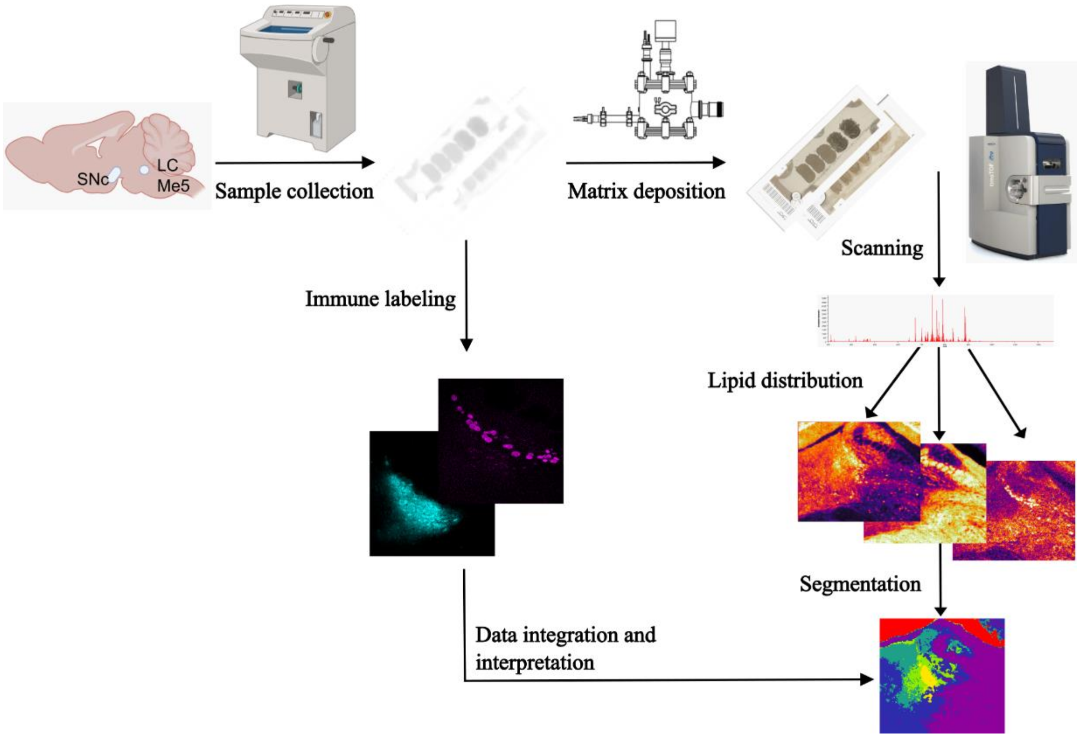


Figure S1. Workflow of the experiment. Sections of male and female mice brain were obtained with the aid of a cryomicrotome. Alternate sections were deposited in plain and ITO-coated microscope slides, so the imaging mass spectrometry (IMS) experiments were accompanied by immunofluorescence (if) images to certify the identity of the neurons. Segmentation was guided by the IF images.

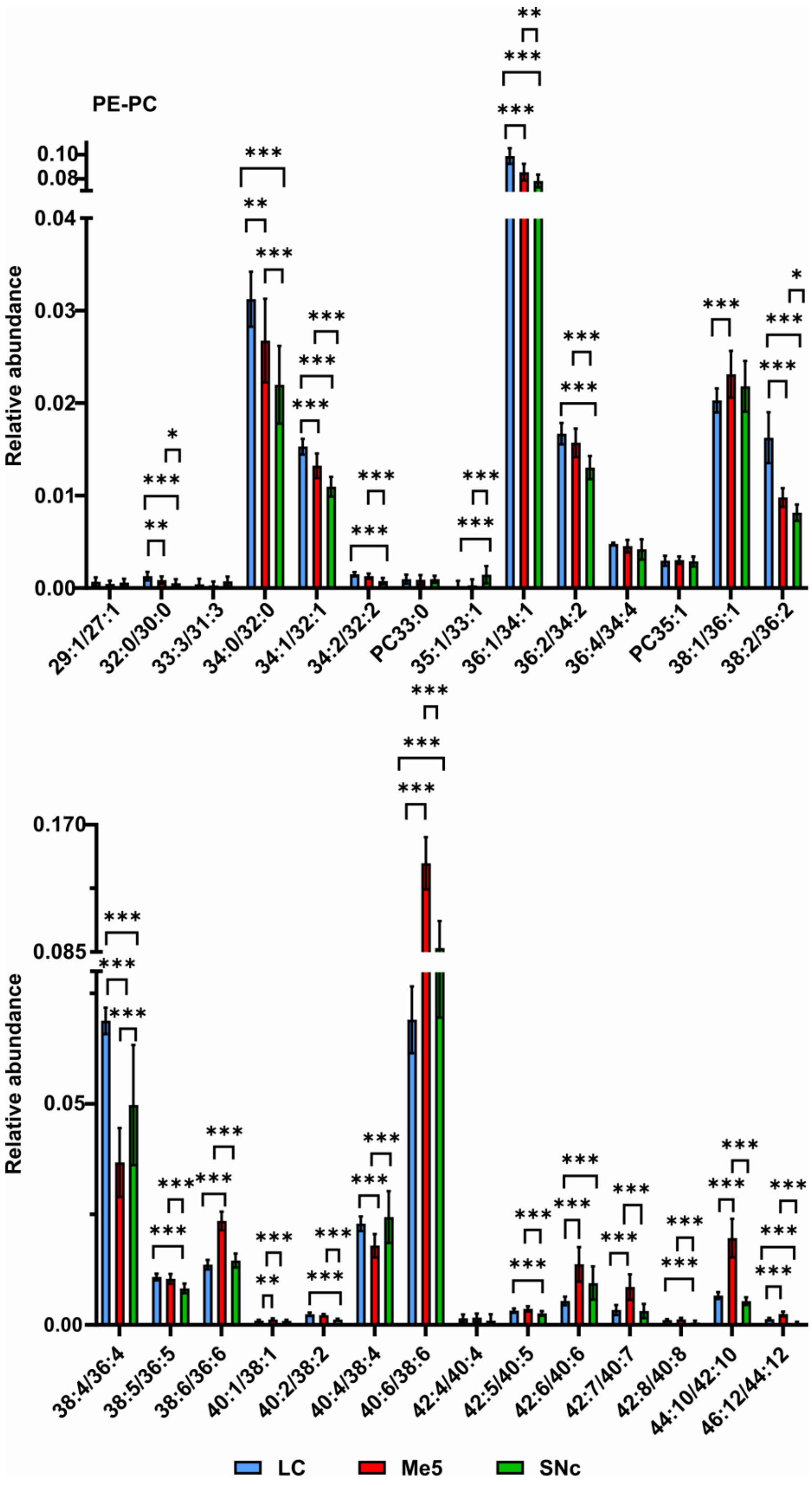


Figure S2. Comparison of the relative abundance of lipid species between neurons of the LC, Me5 and SNc. PE and PC are shown together due to the overlap of several species in the same m/z. For example, 29:1/27:1 corresponds to PE 29:1/PC 27:1. The comparison was divided into two panels for clarity. * = $p<0.05$, ** = $p<0.01$, *** = $p<0.001$

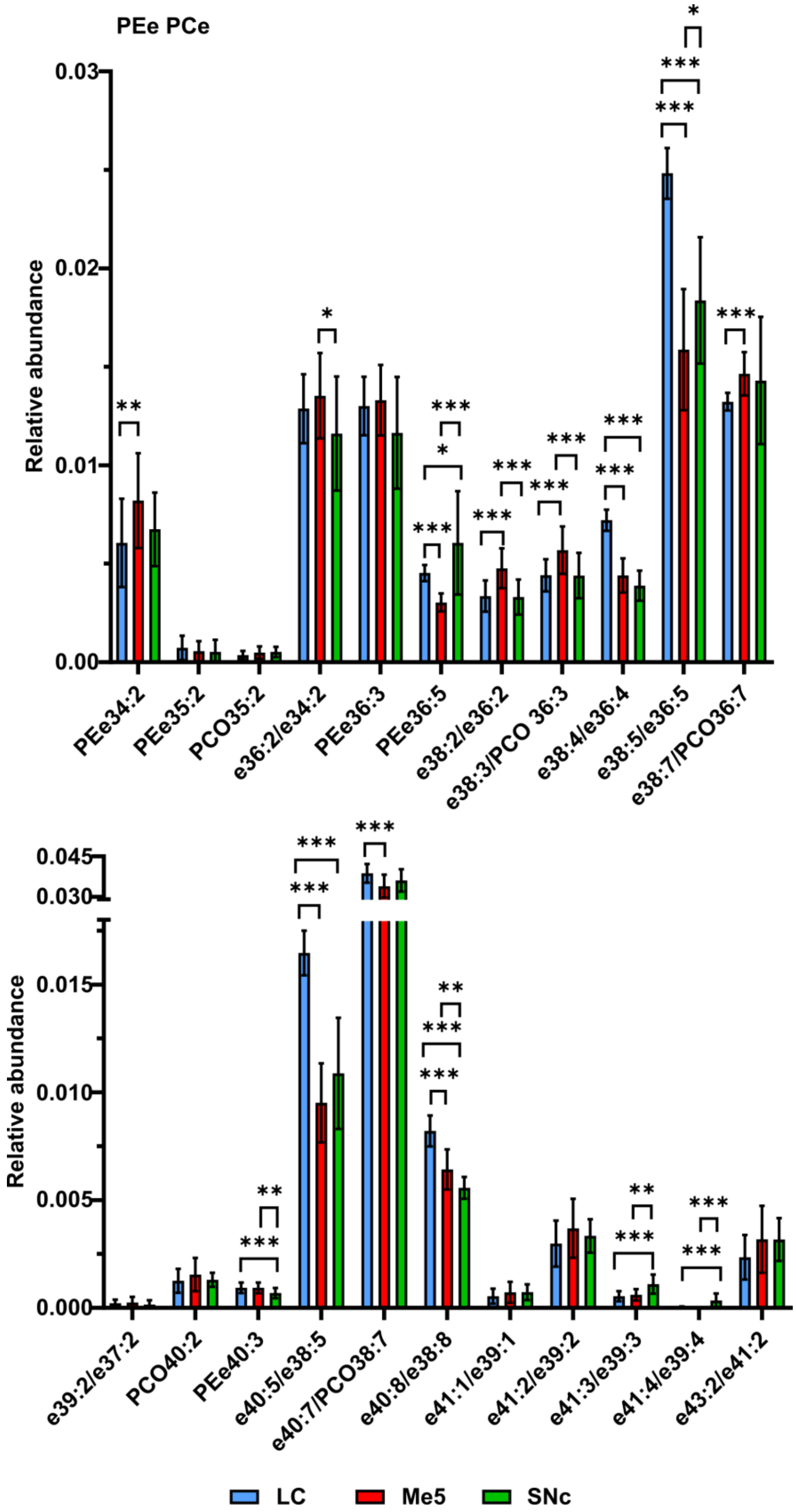


Figure S3. Comparison of the relative abundance of lipid species between neurons of the LC, Me5 and SNc. PEe and PCe are shown together due to the overlap of several species in the same m/z. PEe/PCe was divided into two panels for clarity. * = p<0.05, ** = p<0.01, *** = p<0.001.

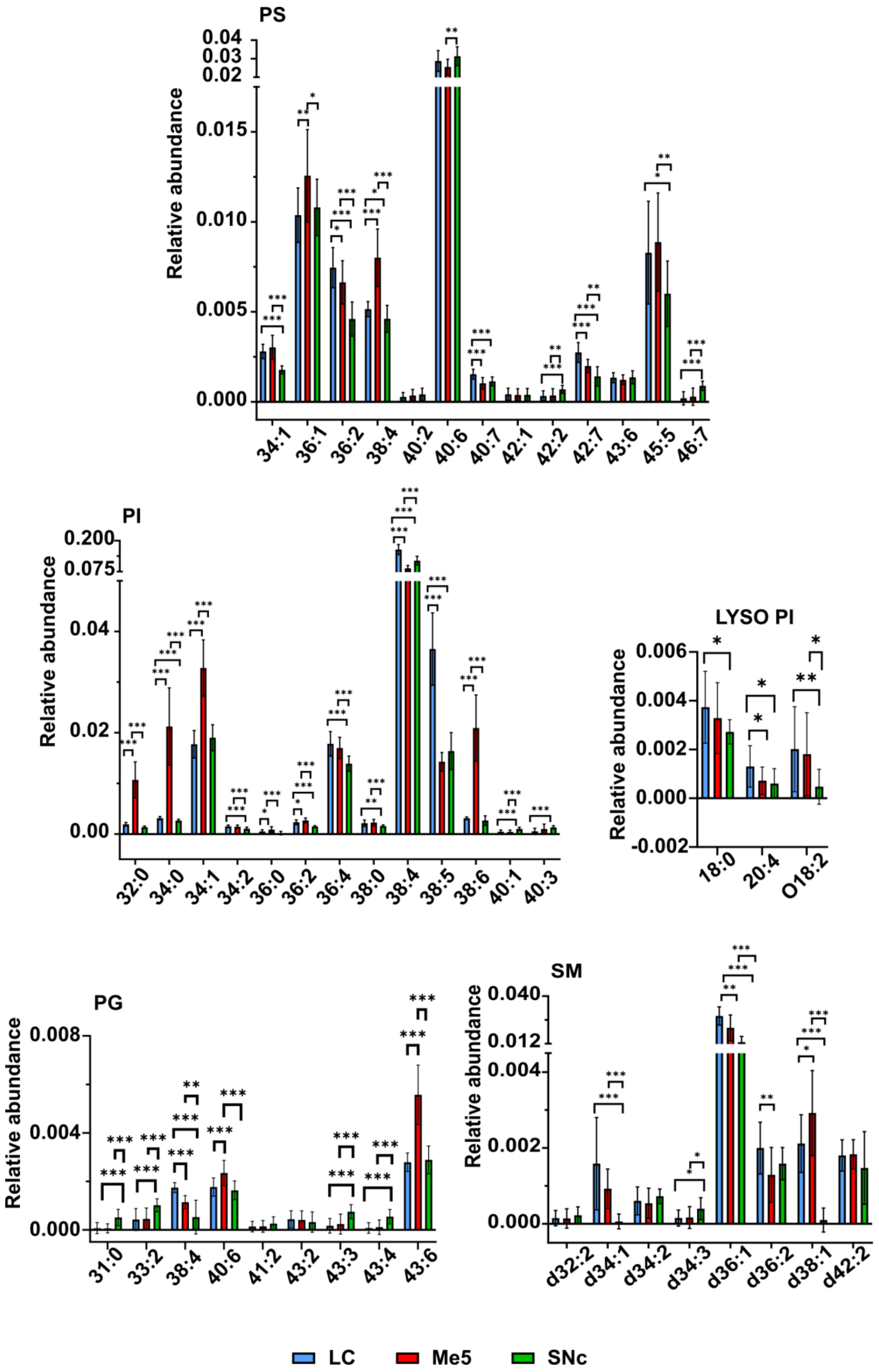


Figure S4. Comparison of the relative abundance of lipid species between neurons of the LC, Me5 and SNc. * = p<0.05, ** = p<0.01, *** = p<0.001.

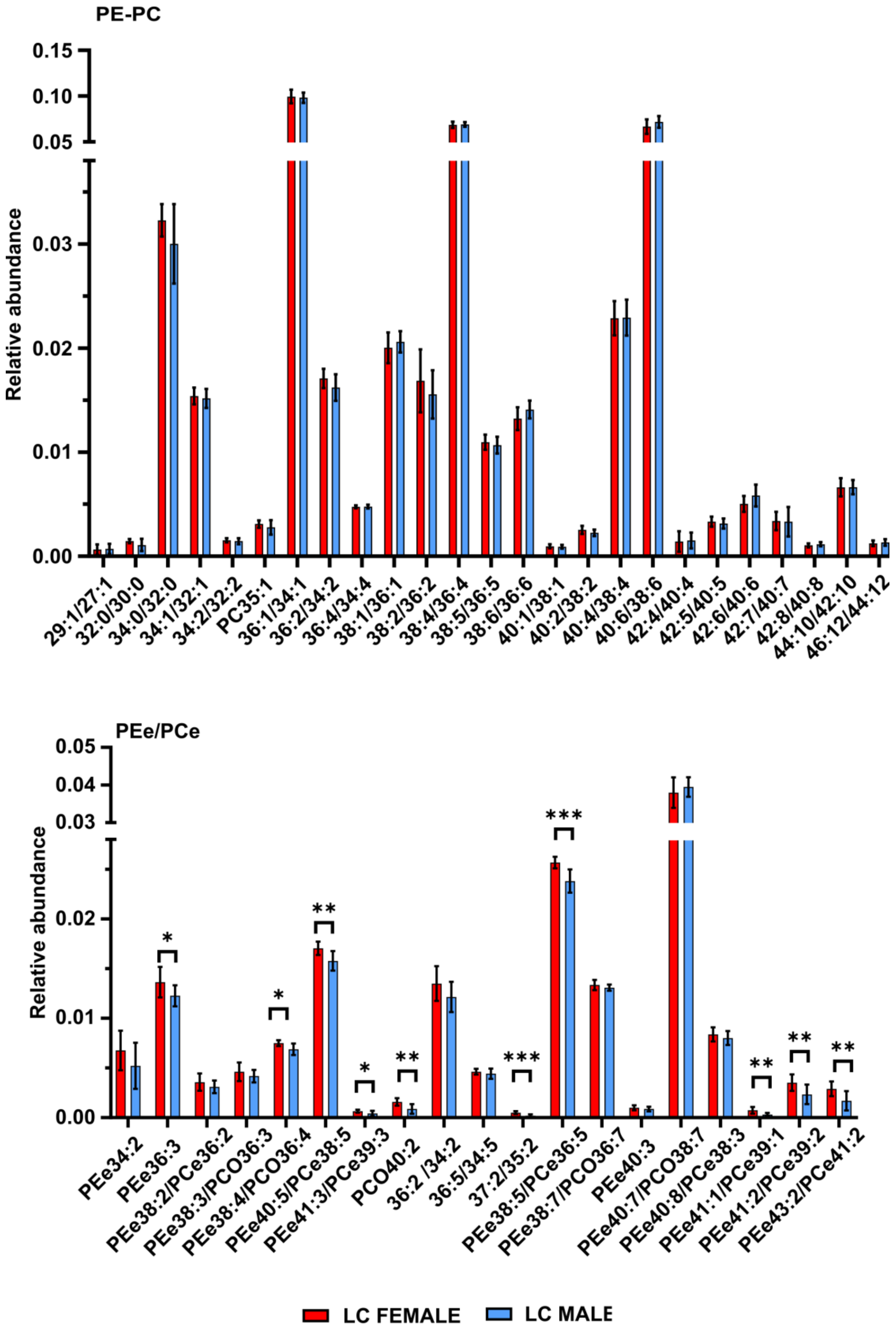


Figure S5. Comparison of the relative abundance of lipid species between LC neurons of male and female mice. PE and PC are shown together due to the overlap of several species in the same m/z. For example, 29:1/27:1 corresponds to PE 29:1/PC 27:1. PEe and PCe were also grouped for the same reason. * = $p<0.05$, ** = $p<0.01$, *** = $p<0.001$.

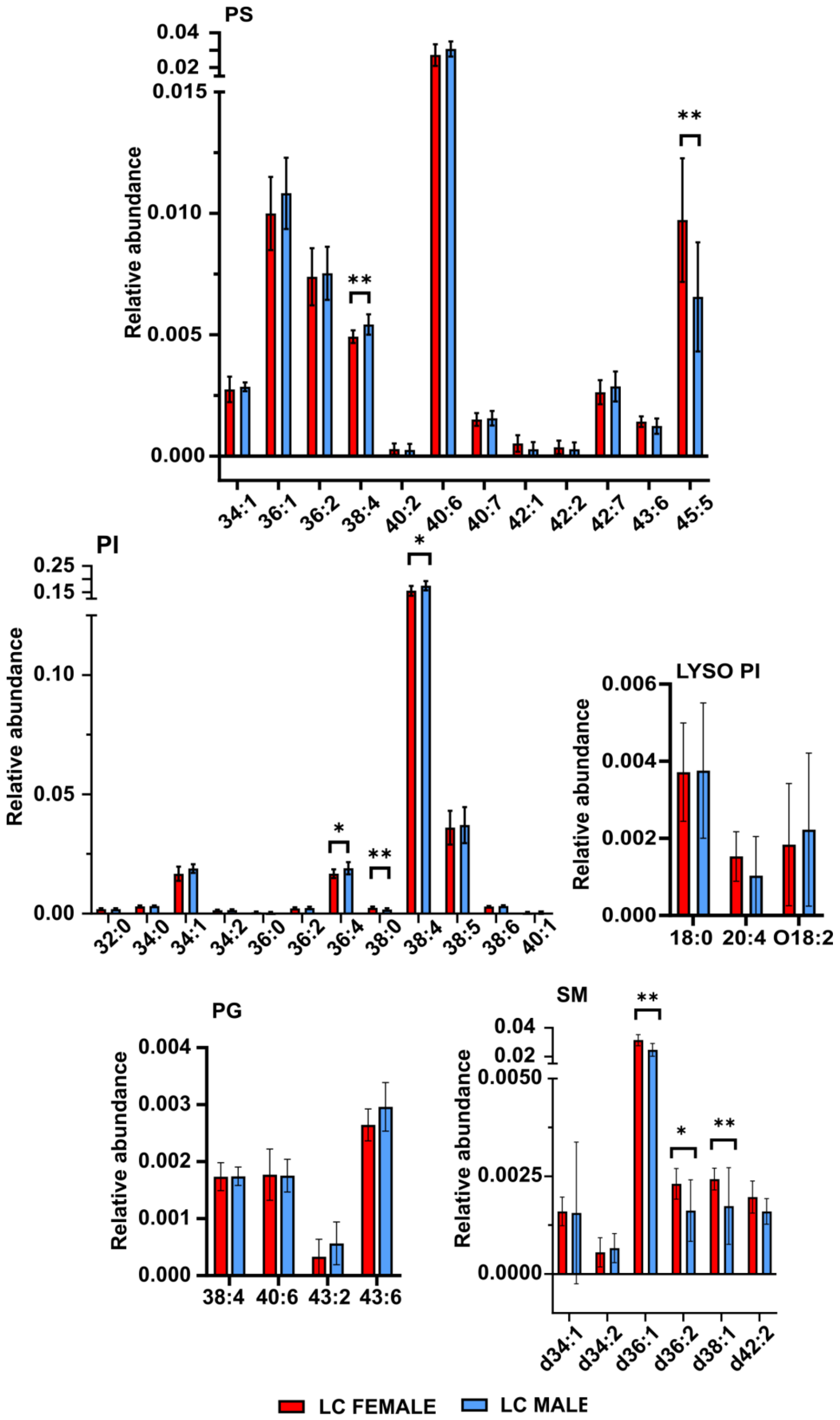


Figure S6. Comparison of the relative abundance of lipid species between LC neurons of male and female mice. For the rest of the classes see Figure S4. * = p<0.05, ** = p<0.01, *** = p<0.001.

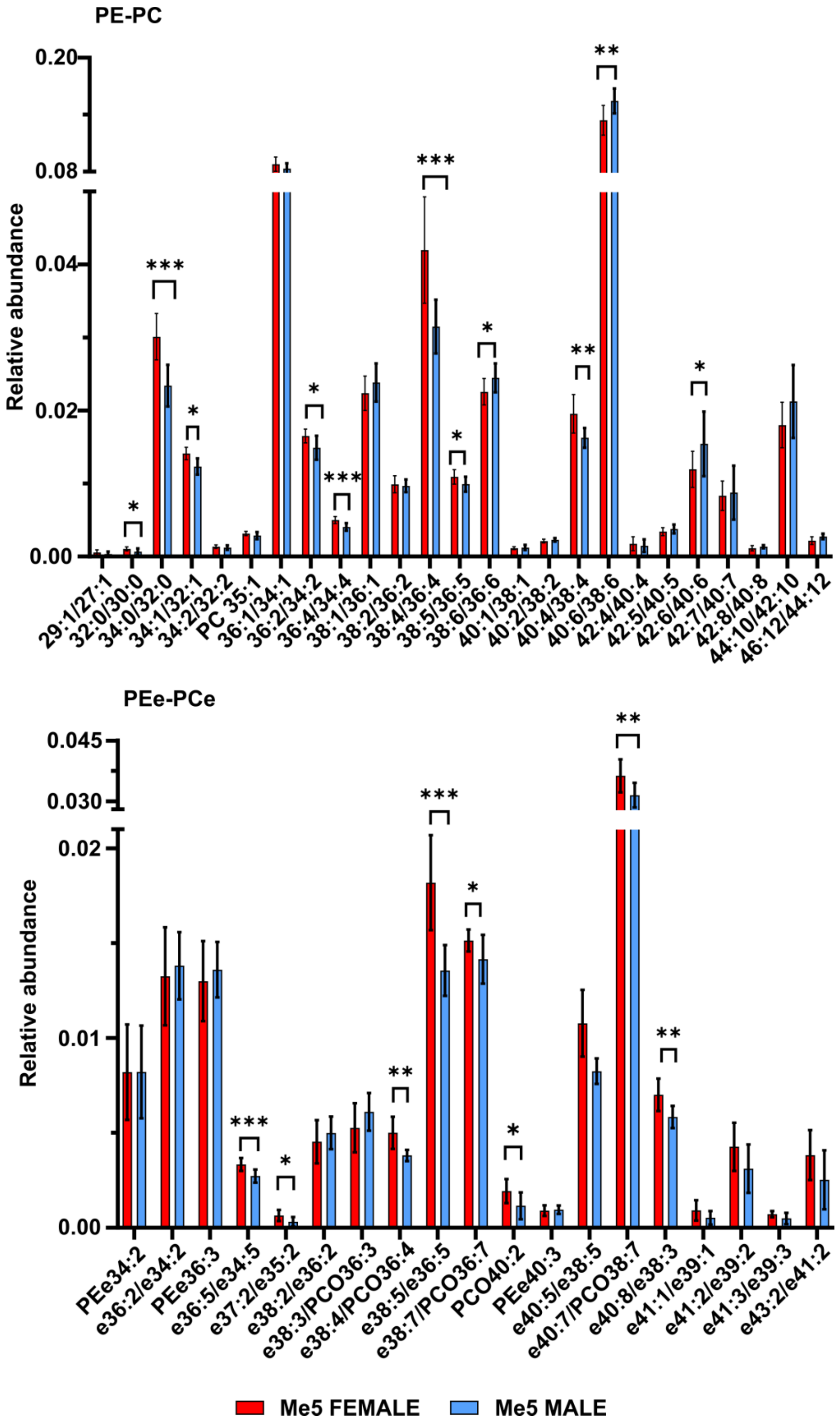


Figure S7. Comparison of the relative abundance of lipid species between Me5 neurons of male and female mice. PE and PC are shown together due to the overlap of several species in the same m/z. For example, 29:1/27:1 corresponds to PE 29:1/PC 27:1. PEe and PCe were also grouped for the same reason. * = p<0.05, ** = p<0.01, *** = p<0.001.

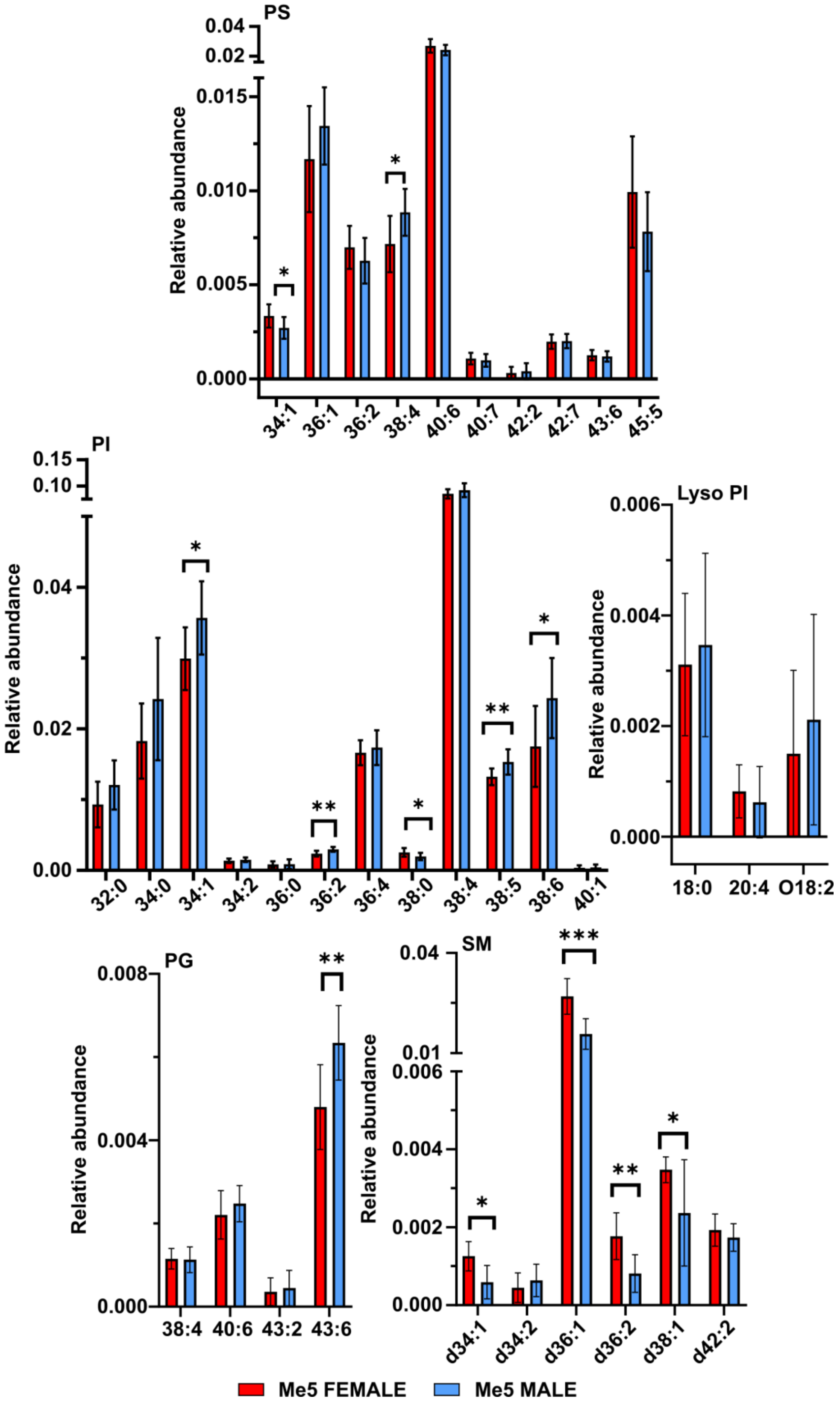


Figure S8. Comparison of the relative abundance of lipid species between Me5 neurons of male and female mice. For the rest of the classes see Figure S6. * = $p<0.05$, ** = $p<0.01$, *** = $p<0.001$.

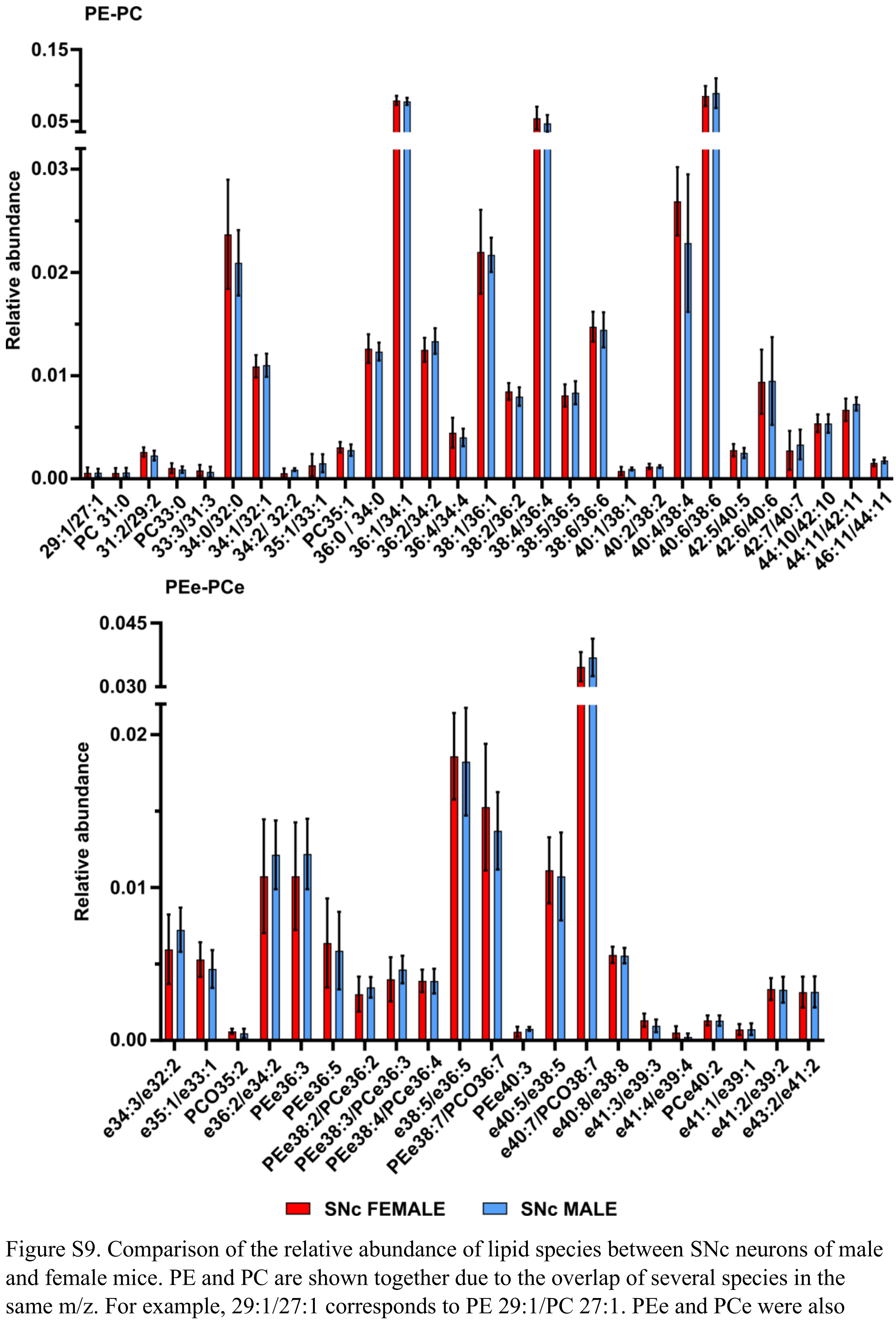


Figure S9. Comparison of the relative abundance of lipid species between SNc neurons of male and female mice. PE and PC are shown together due to the overlap of several species in the same m/z. For example, 29:1/27:1 corresponds to PE 29:1/PC 27:1. PEe and PCe were also grouped for the same reason. For the rest of the classes see Figure S9. * = p<0.05, ** = p<0.01, *** = p<0.001.

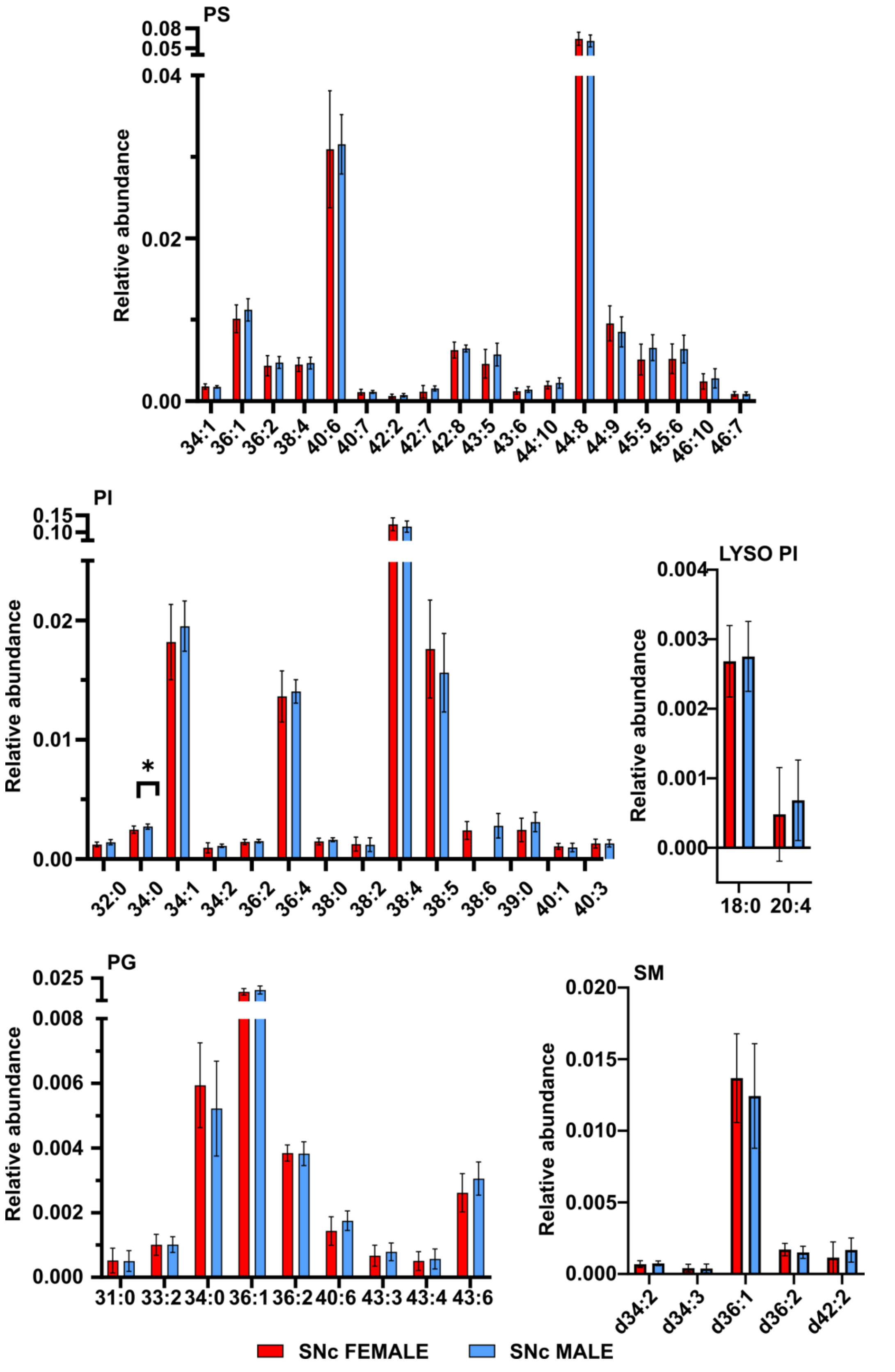


Figure S10. Comparison of the relative abundance of lipid species between MSN neurons of male and female mice. For the rest of the classes see Figure S8. * = p<0.05, ** = p<0.01, *** = p<0.001.
